## Supplementary files for "A benchmark study of simulation methods for single-cell RNA sequencing data"

Supplementary Figure

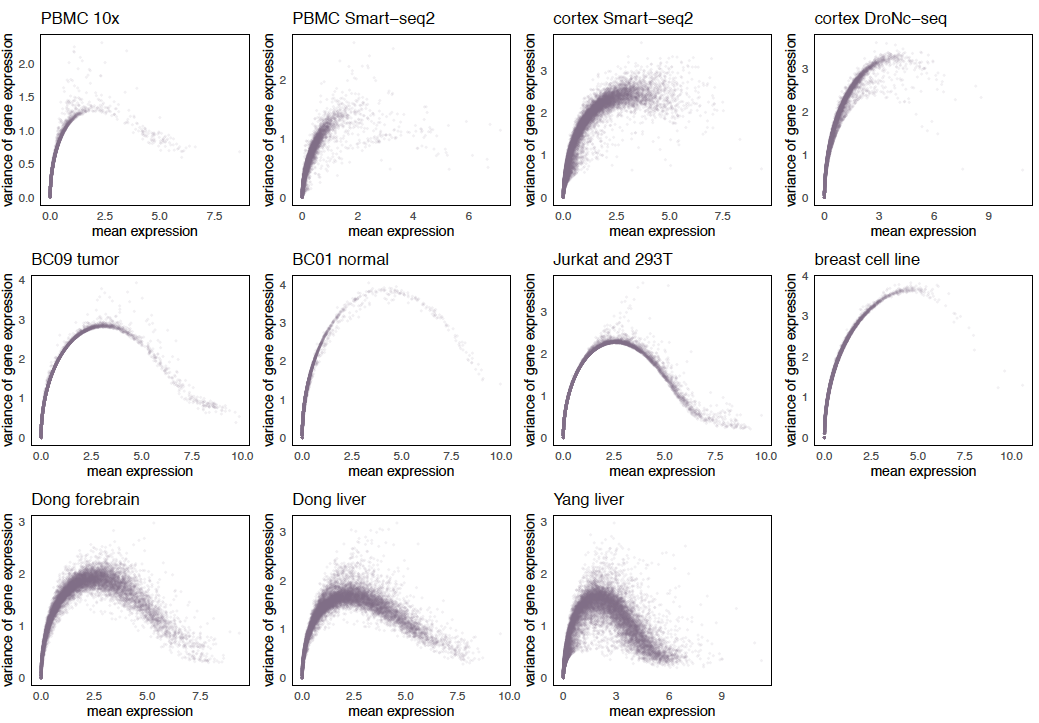

### **Supplementary Figure 1. Mean-variance relationship of genes across multiple datasets.**

We show the property of mean-variance relationship of genes in 11 datasets as an example to demonstrate the variability of data properties across datasets. Top panel shows four datasets of different protocols, the first two from human PBMC samples and the latter two from mouse cortex samples. Middle panel shows two datasets from tissue source and two datasets with cell line source. Bottom panel shows datasets of multiple cell types in mouse sample.

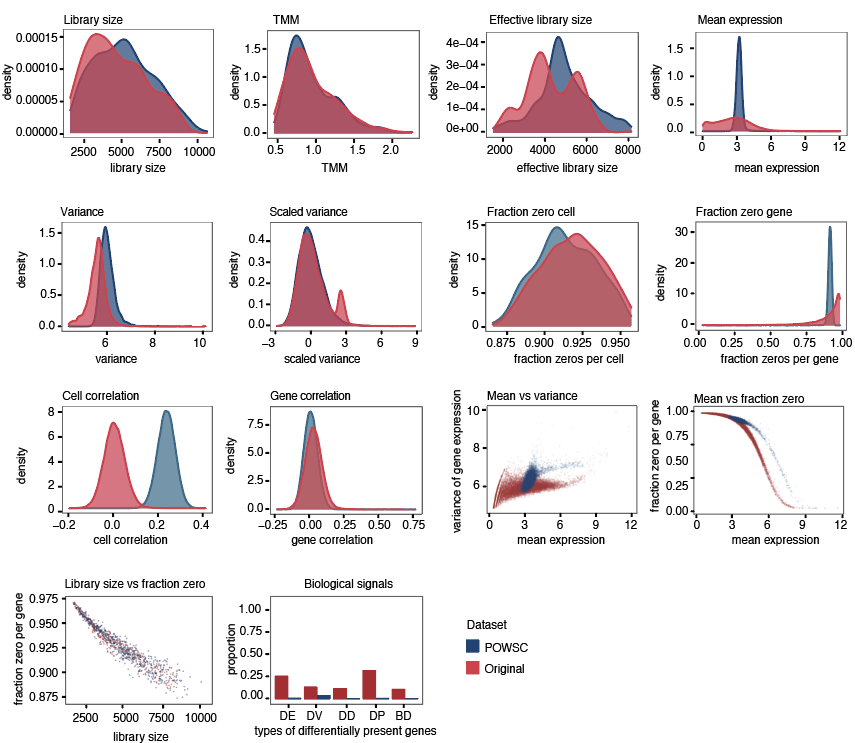

### **Supplementary Figure 2. Visual representation of the evaluation criteria in properties estimation and biological signals.**

As an illustrative example, we compared the simulation data generated by POWSC and the original dataset Soumillon that was used as the reference input. In properties estimation, we compared the concordance of the data characteristics across multiple properties using the KDE statistic. In biological signals, we compared the concordance of the amount of proportion of biological signals in simulated and in real data.

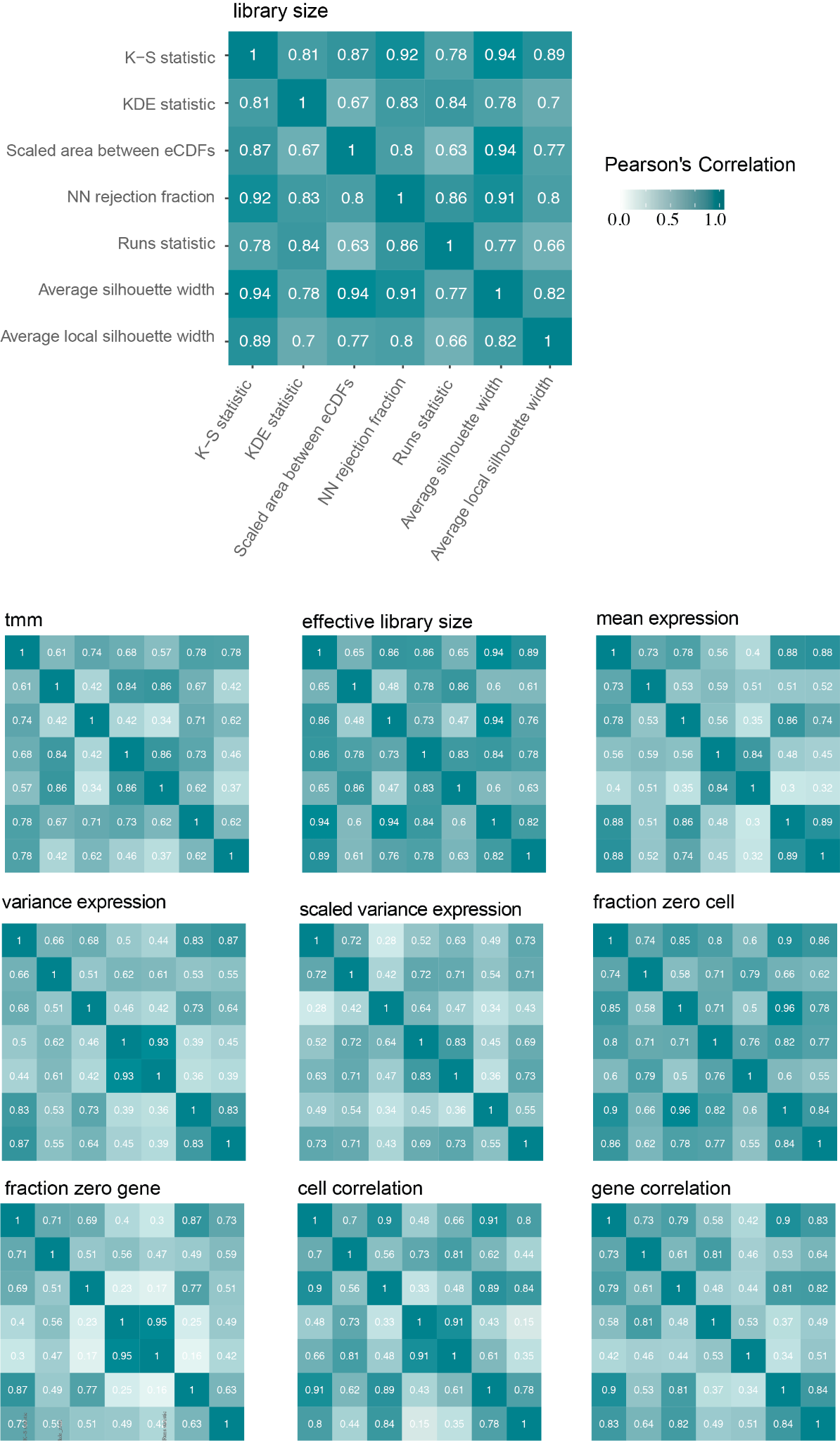

### **Supplementary Figure 3. Correlation between seven measures on quantifying similarities for univariate properties.**

Top panel shows the correlation matrix for the property library size, enlarged for readability of axis labels. Bottom panel shows correlation matrix for the remaining univariate properties. The axis labels are consistent and are not shown for readability of the matrix.

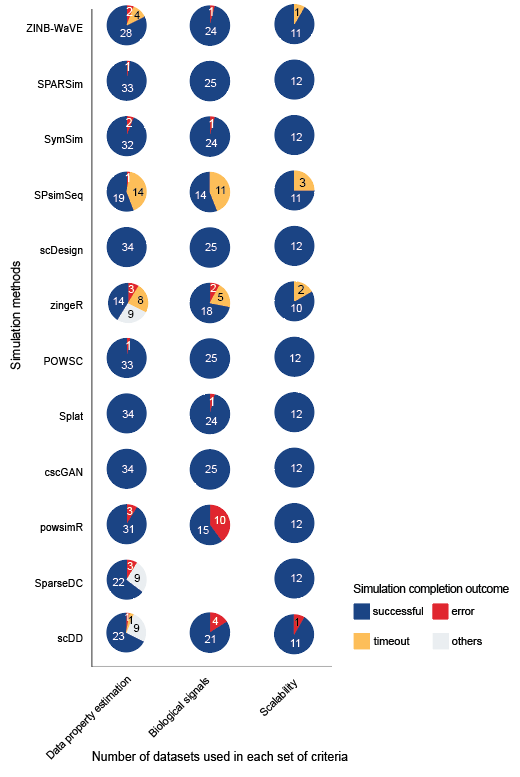

**Supplementary Figure 4. Number of datasets used for evaluation and simulation completion outcome**

A total of 34 datasets were used for evaluating the simulation methods on data property estimation, 25 datasets were used for evaluating biological signals and 12 datasets were used for evaluating scalability (see Methods). Pie chart denotes the completion outcome of each simulation method. “Successful” indicates the method produced a simulation dataset within the given time limit. “Error” indicates the method encountered issues during simulation. “Timeout” indicates the method was not able to finish the simulation within the given time limit (see Methods). “Others” indicates a special situation where the methods require the dataset to have two or more cell types and therefore could not be tested on datasets containing a single cell type.

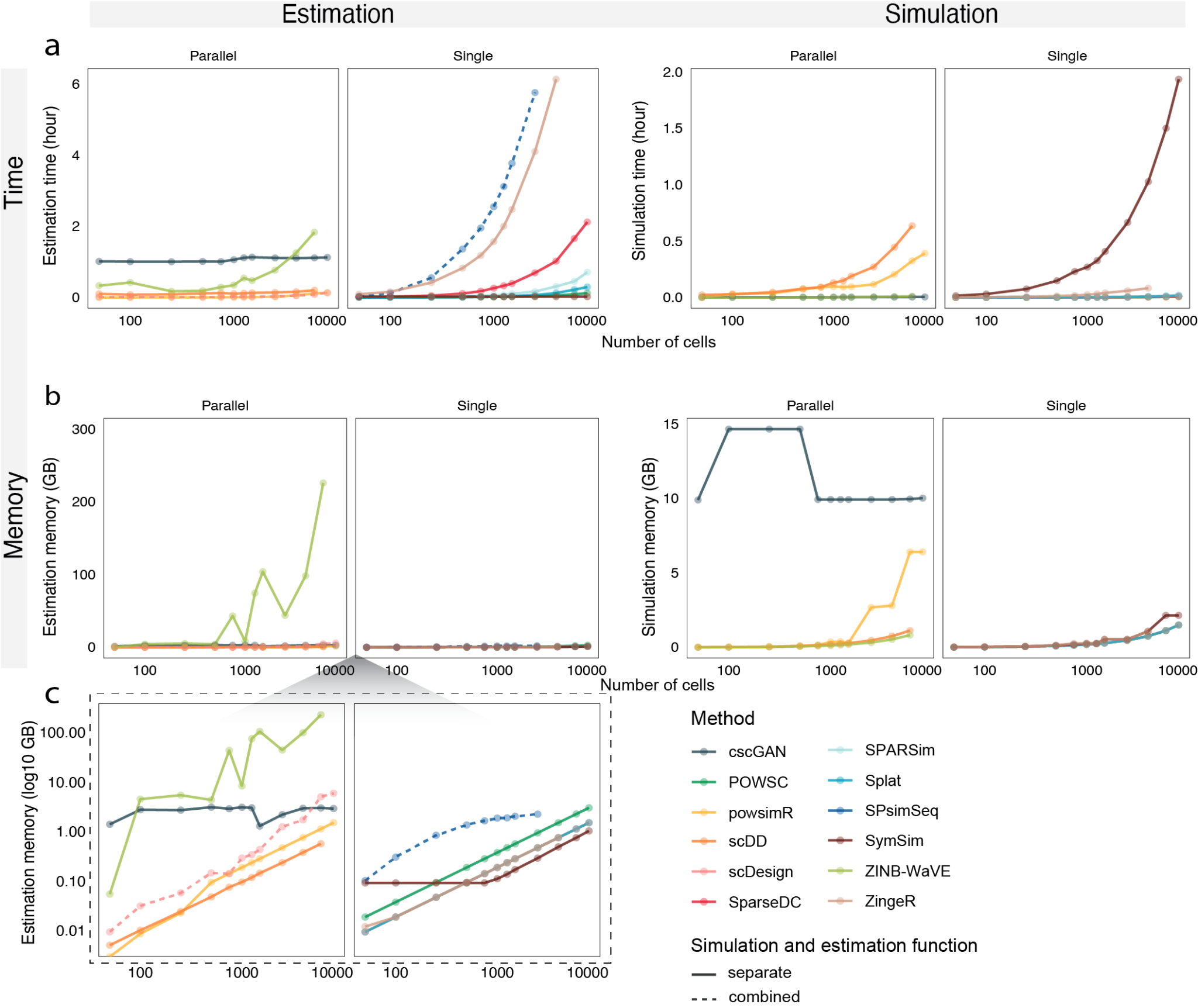

### **Supplementary Figure 5. Run time and memory consumption of each method.**

**a** Runtime of each method. **b** Maximal memory usage of each method. The number of cells is shown in log10 scale. Methods that support parallel computing and those that only support single core are shown separately. Most methods involve a two-step process of properties estimation and dataset simulation. For those methods, we recorded and shown results for the two steps separately under the estimation and simulation panels. A solid line was used to indicate these methods. For methods that perform the two steps together in a single function, we displayed the results under the estimation panel. A dashed line was used to indicate these methods. **c** This shows the same result as in **b**, but with the y-axis in log10 scale for enhanced readability.

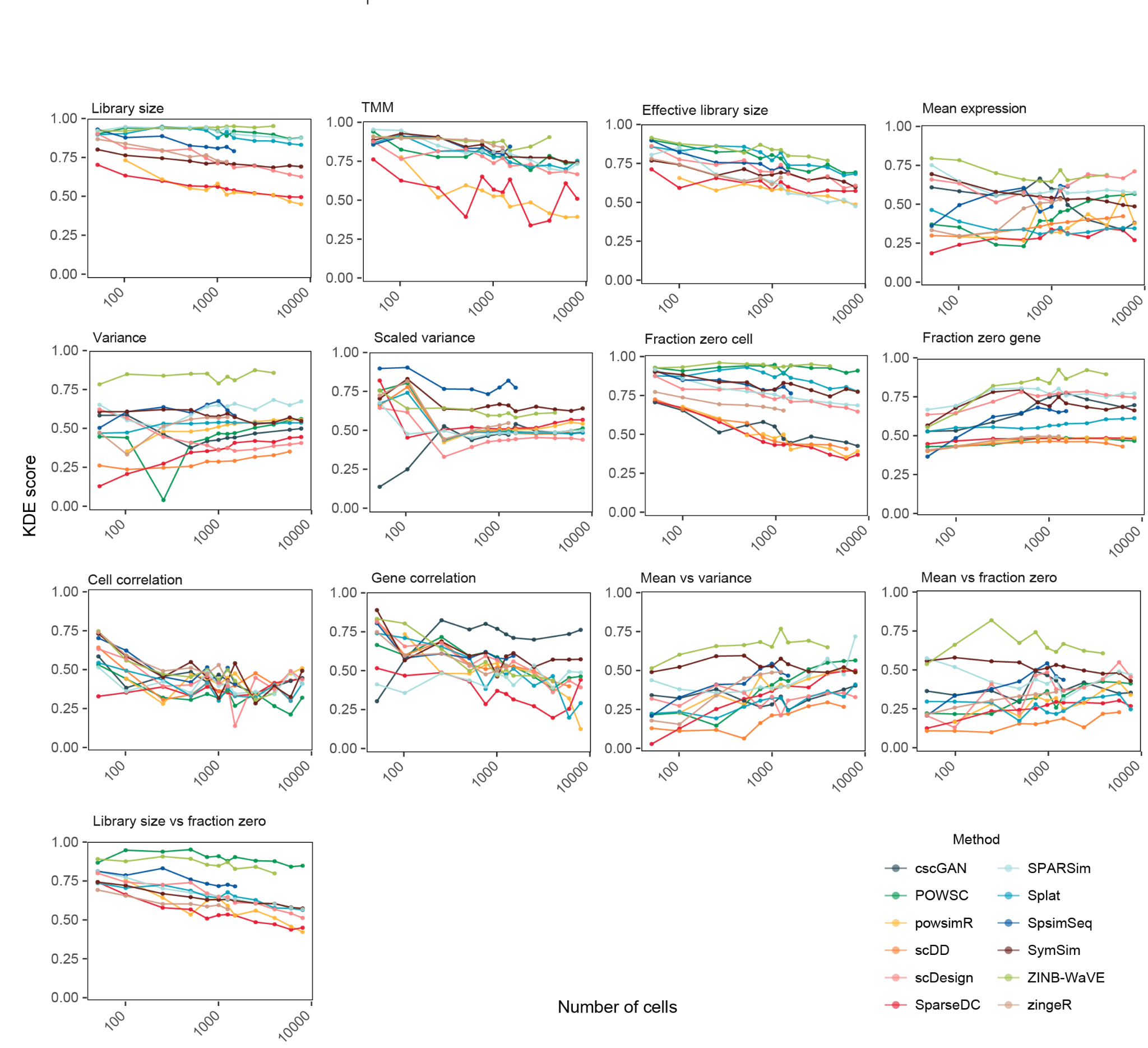

### **Supplementary Figure 6. Impact of the number of cells on property estimation.**

The x-axis shows the number of cells in log10 scale and y-axis shows the score. The line shows the trends with increasing cell numbers. The dot indicates where a measurement is taken. Each measurement was taken three times and the average was shown in the figure.

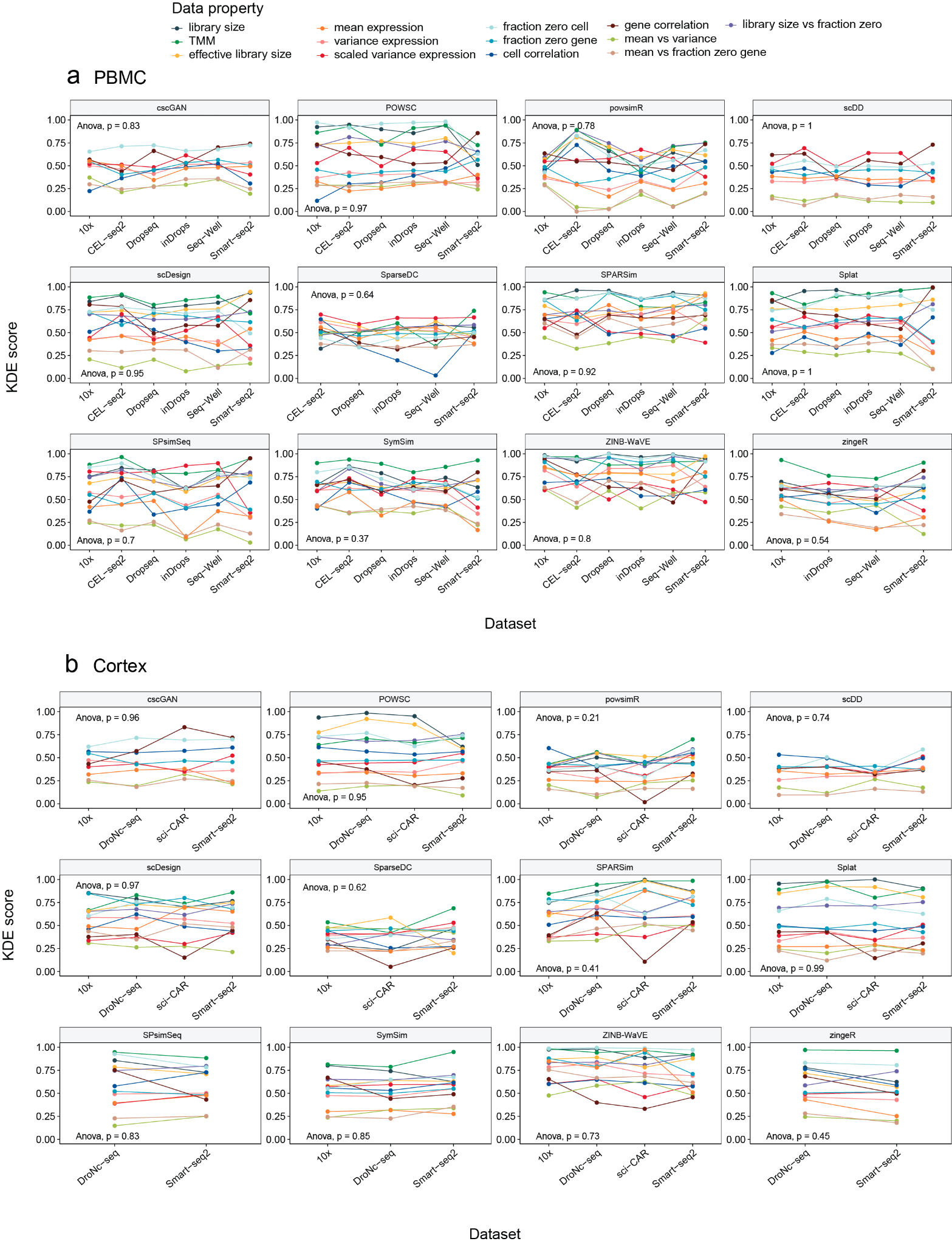

### **Supplementary Figure 7. Impact of sequencing protocols on data property estimation.**

The impact of sequencing protocols on data property estimation using **a** human PBMC data collections and **b** mouse cortex data collections. ANOVA was performed to examine the statistical significance of the change in KDE score due to sequencing protocol effect.

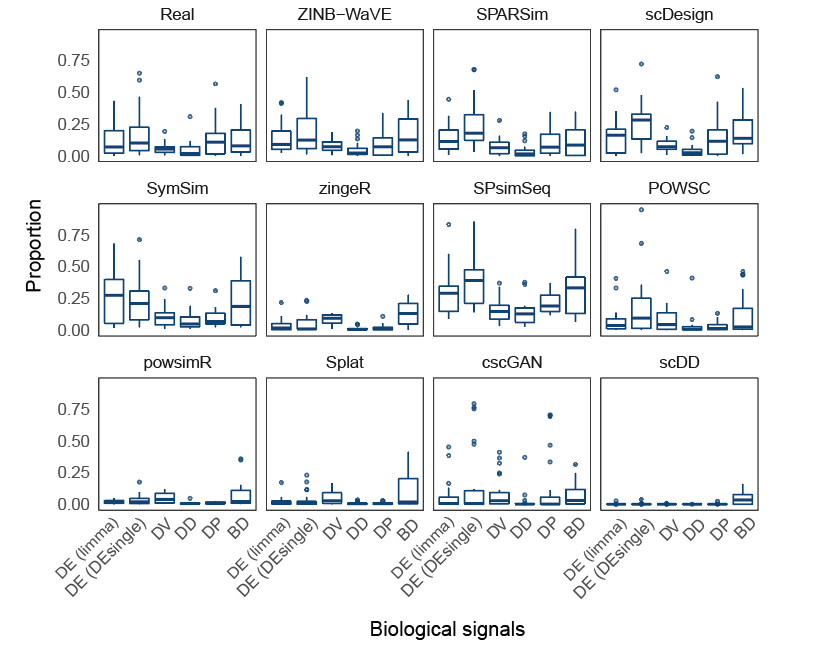
**Supplementary Figure 8. Proportion of biological signals in real and simulated data generated by simulation methods.**

The boxplots show the distribution of the proportion of biological signals for all datasets examined. The proportion of biological signals in the simulated data ideally should be similar to that of the real data.

Supplementary Table

**Supplementary Table 1. Additional details of the scRNA-seq simulation methods evaluated in this study.**

| **Methods** | **Implementation language** | **Year of publication** | **Reference (doi)** | **Software version** | **Input data (raw/**  **normalised)** | **Output data (raw**  **/normalised)** |
| --- | --- | --- | --- | --- | --- | --- |
| scDD | R | 2016 | 10.1186/s13059-016-1077-y | 1.12.0 (implemented in Splatter) | raw | normalised |
| Splat | R | 2017 | 10.1186/s13059-017-1305-0 | 1.12.0 | raw | raw |
| powsimR | R | 2017 | 10.1093/bioinformatics/btx435 | 1.2.3 | raw | raw |
| SparseDC | R | 2017 | 10.1093/nar/gkx1113 | 0.1.17 (implemented in Splatter) | raw | raw |
| zingeR | R | 2018 | 10.1186/s13059-018-1406-4 | 0.1.0 | raw | raw |
| ZINB-WaVE | R | 2018 | 10.1038/s41467-017-02554-5 | 1.10.0 (implemented in Splatter) | raw | raw |
| SymSim | R | 2019 | 10.1038/s41467-019-10500-w | 0.0.0.9000 | raw | raw |
| scDesign | R | 2019 | 10.1093/bioinformatics/btz321 | 1.0.0 | raw | raw |
| SPARSim | R | 2020 | 10.1093/bioinformatics/btz752 | 0.9.5 | both raw and normalised | raw |
| SPsimSeq | R | 2020 | 10.1093/bioinformatics/btaa105 | 0.99.13 | raw | raw |
| POWSC | R | 2020 | 10.1093/bioinformatics/btaa607 | 0.1.0 | raw | raw |
| cscGAN | Python | 2020 | 10.1038/s41467-019-14018-z | GitHub version 379ff6e | raw | normalised |

**Supplementary Table 2. Details of the datasets used in this study.**

| **Dataset** | **Accession** | **Name** | **Description** | **Species** | **Protocol** | **Number of cells** | **Multiple cell types/**  **condition ?** | **Source** |
| --- | --- | --- | --- | --- | --- | --- | --- | --- |
| 1 | SCP425 | cortex sciRNAseq | Comparison of four protocols using mouse cortex | Mouse | sci-RNA-seq | 4912 | Yes | https://singlecell.broadinstitute.org/single_cell/study/SCP425/single-cell-comparison-cortex-data#study-download |
| 2 | SCP425 | cortex 10x |  | Mouse | 10x Genomics | 5367 | Yes | https://singlecell.broadinstitute.org/single_cell/study/SCP425/single-cell-comparison-cortex-data#study-download |
| 3 | SCP425 | cortex DroNc-seq |  | Mouse | DroNc-seq | 2345 | Yes | https://singlecell.broadinstitute.org/single_cell/study/SCP425/single-cell-comparison-cortex-data#study-download |
| 4 | SCP425 | cortex Smart-seq2 |  | Mouse | Smart-seq2 | 644 | Yes | https://singlecell.broadinstitute.org/single_cell/study/SCP425/single-cell-comparison-cortex-data#study-download |
| 5 | SCP424 | PBMC 10x | Comparison of six protocols using Human Peripheral Blood Mononuclear Cell | Human | 10x Genomics | 3312 | Yes | https://singlecell.broadinstitute.org/single_cell/study/SCP424/single-cell-comparison-pbmc-data#study-summary |
| 6 | SCP424 | PBMC CEL-seq2 |  | Human | CEL-seq2 | 526 | Yes | https://singlecell.broadinstitute.org/single_cell/study/SCP424/single-cell-comparison-pbmc-data#study-summary |
| 7 | SCP424 | PBMC Drop-seq |  | Human | Drop-seq | 6357 | Yes | https://singlecell.broadinstitute.org/single_cell/study/SCP424/single-cell-comparison-pbmc-data#study-summary |
| 8 | SCP424 | PBMC inDrops |  | Human | inDrops | 4184 | Yes | https://singlecell.broadinstitute.org/single_cell/study/SCP424/single-cell-comparison-pbmc-data#study-summary |
| 9 | SCP424 | PBMC Seq-Well |  | Human | Seq-Well | 2908 | Yes | https://singlecell.broadinstitute.org/single_cell/study/SCP424/single-cell-comparison-pbmc-data#study-summary |
| 10 | SCP424 | PBMC Smart-seq2 |  | Human | Smart-seq2 | 522 | Yes | https://singlecell.broadinstitute.org/single_cell/study/SCP424/single-cell-comparison-pbmc-data#study-summary |
| 11 | see source | Tabula Muris | The 10x subset of Tabula Muris | Mouse | 10x Genomics | 55656 | Yes | https://tabula-muris.ds.czbiohub.org/ |
| 12 | GSE114724 | BC09 tumor | Tumor of breast cancer patient ID BC09 | Human | 10x Genomics | 7000 | No | https://www.ncbi.nlm.nih.gov/geo/query/acc.cgi?acc=GSE114724 |
| 13 | GSE114725 | BC02 tumor | Tumor of breast cancer patient ID BC02 | Human | inDrops | 2437 | No | https://www.ncbi.nlm.nih.gov/geo/query/acc.cgi?acc=GSE114725 |
| 14 | GSE114725 | BC01 blood | Blood of breast cancer patient ID BC01 | Human | inDrops | 3034 | No | https://www.ncbi.nlm.nih.gov/geo/query/acc.cgi?acc=GSE114725 |
| 15 | GSE114725 | BC02 lymph | Lymph node of breast cancer patient ID BC02 | Human | inDrops | 6129 | No | https://www.ncbi.nlm.nih.gov/geo/query/acc.cgi?acc=GSE114725 |
| 16 | GSE114725 | BC01 normal | Normal breast tissue of breast cancer patient ID BC01 | Human | inDrops | 3607 | No | https://www.ncbi.nlm.nih.gov/geo/query/acc.cgi?acc=GSE114725 |
| 17 | GSE106202 | breast cell line | MDA-MB-231 cells cultured in glucose | Human | Drop-seq | 785 | No | https://www.ncbi.nlm.nih.gov/geo/query/acc.cgi?acc=GSE106202 |
| 18 | GSE102827 | light endo | Endothelial smooth muscle of primary visual cortex from mice, exposed to light for 0h, 1h and 4h | Mouse | inDrops | 4071 | Yes | https://www.ncbi.nlm.nih.gov/geo/query/acc.cgi?acc=GSE102827 |
| 19 | GSE102827 | light micro | Microglia of primary visual cortex from visually stimulated mice, exposed to light for 0h, 1h and 4h | Mouse | inDrops | 10158 | Yes | https://www.ncbi.nlm.nih.gov/geo/query/acc.cgi?acc=GSE102827 |
| 20 | GSE92495 | Gierahn | human HEK293 (embryonic kidney cells) cell line | Human | Seq-Well | 1453 | No | https://www.ncbi.nlm.nih.gov/geo/query/acc.cgi?acc=GSE92495 |
| 21 | see source | 293T | 293T (adenovirus-immortalized human embryonic kidney cells) cell line | Human | 10x Genomics | 2885 | No | https://support.10xgenomics.com/single-cell-gene-expression/datasets/1.1.0/293t |
| 22 | see source | Jurkat and 293T | mixture of Jurkat (human T lymphocyte) and 293T | Human | 10x Genomics | 6143 | Yes | https://support.10xgenomics.com/single-cell-gene-expression/datasets/1.1.0/jurkat |
| 23 | GSE77288 | Tung | Three iPSC ( Induced Pluripotent Stem Cells) lines | Human | SMARTer | 564 | Yes | https://www.ncbi.nlm.nih.gov/geo/query/acc.cgi?acc=GSE77288 |
| 24 | GSE113660 | Chen | Rh41(human alveolar rhabdomyosarcoma) cell line | Human | 10x Genomics | 6875 | Yes | https://www.ncbi.nlm.nih.gov/geo/query/acc.cgi?acc=GSE113660 |
| 25 | GSE60361 | Zeisel | cortex of mice | Mouse | STRT-seq | 3005 | Yes | https://www.ncbi.nlm.nih.gov/geo/query/acc.cgi?acc=GSE60361 |
| 26 | GSE72857 | Pual | Bone marrow myeloid progenitors | Mouse | MARS-seq | 6144 | Yes | https://www.ncbi.nlm.nih.gov/geo/query/acc.cgi?acc=GSE72857 |
| 27 | GSE63472 | retina | Mouse retina | Mouse | Drop-seq | 6598 | No | https://www.ncbi.nlm.nih.gov/geo/query/acc.cgi?acc=GSE63472 |
| 28 | GSE87038 | Dong forebrain | Forebrain cells of E9.5 to E11.5 mouse embryos | Mouse | Smart-seq2 | 196 | Yes | https://www.ncbi.nlm.nih.gov/geo/query/acc.cgi?acc=GSE87038 |
| 29 | GSE87038 | Dong skin | Skin cells of E9.5 to E11.5 mouse embryos | Mouse | Smart-seq2 | 196 | Yes | https://www.ncbi.nlm.nih.gov/geo/query/acc.cgi?acc=GSE87038 |
| 30 | GSE87038 | Dong intest | Intestine cells of E9.5 to E11.5 mouse embryos | Mouse | Smart-seq2 | 196 | Yes | https://www.ncbi.nlm.nih.gov/geo/query/acc.cgi?acc=GSE87038 |
| 31 | GSE87038 | Dong liver | Liver cells of E9.5 to E11.5 mouse embryos | Mouse | Smart-seq2 | 196 | Yes | https://www.ncbi.nlm.nih.gov/geo/query/acc.cgi?acc=GSE87038 |
| 32 | GSE90047 | Yang liver | Liver cells of E10.5 to E17.5 mouse embryos | Mouse | Smart-seq2 | 447 | Yes | https://www.ncbi.nlm.nih.gov/geo/query/acc.cgi?acc=GSE90047 |
| 33 | GSE75748 | stem cell | Human pluripotent stem cells (hPSCs) | Human | SMARTer | 758 | Yes | https://www.ncbi.nlm.nih.gov/geo/query/acc.cgi?acc=GSE75748 |
| 34 | GSE112004 | Francesconi | B cell precursors from bone marrow, induced to either trans-differentiate to macrophages or to reprogram into iPSCs | Mouse | MARS-Seq | 3833 | Yes | https://www.ncbi.nlm.nih.gov/geo/query/acc.cgi?acc=GSE112004 |
| 35 | GSE53638 | Soumillon | differentiating cells of human adipose-derived stem/stromal cells | Human | SCRB-Seq | 2968 | Yes | https://www.ncbi.nlm.nih.gov/geo/query/acc.cgi?acc=GSE53638 |

**Supplementary Table 3. Detailed simulation strategy of each method.**

| **Methods** | **Simulation Strategy for evaluating data property estimation** | **Simulation Strategy for evaluating biological signals** |
| --- | --- | --- |
| Splat | Estimated the parameters and simulated each cell type separately. | Estimated parameters from the largest cell type in a dataset, set the number of groups to 2 and the proportion of differential expressed (DE) genes to the proportion between the two largest cell types in the dataset (*). This is because the genes in the simulated data do not have a one-to-one matching relationship with the input data and hence it is not possible to combine two simulated data generated from two cell types separately. |
| powsimR | This method generates DE genes from a homogenous population, for example, a particular cell type from one patient to create two artificial populations. We therefore estimated the parameters and simulated each cell type separately. The proportion of DE and log fold change were set to be a null scenario to maintain the biological signals in the original cell type population. | This method generates DE genes from a homogenous population. We therefore estimated the parameters and simulated the largest cell type. The proportion of DE was set to the proportion between the two largest cell types in the dataset. |
| SymSim | Estimated the parameters and simulated each cell type separately. | Estimated the parameters and simulated the two largest cell types separately. |
| scDesign | Estimated the parameters and simulated each cell type separately. | This method generates DE genes from a homogenous population. We therefore estimated the parameters and simulated the largest cell type. The proportion of DE was set to the proportion between the two largest cell types in the dataset |
| SPARSim | Estimated the parameters and simulated each cell type separately. | Estimated the parameters and simulated the two largest cell types separately. This is because the method returns gene names in the simulated data and therefore we can combine the two datasets and evaluate the biological signals between the two cell types. |
| SPsimSeq | Estimated the parameters and simulated each cell type separately. | Estimated the parameters and simulated the two largest cell types separately. |
| POWSC | Estimated the parameters and simulated each cell type separately. | Estimated the parameters and simulated the two largest cell types separately. |
| zingeR | We estimated and simulated every two cell types at a time with the proportion of DE gene set to 10%. This is the setting used by the authors of this method when comparing their simulated dataset to the original dataset. | We estimated and simulated the two largest cell types at a time with the proportion of DE gene set to the proportion between these two cell types. |
| scDD | We estimated and simulated every two cell types at a time with the proportion of DE genes set to 10%. This is because the method requires two cell types to be simulated at once with a given proportion of DE genes between them. | We estimated and simulated the largest two cell types with the proportion of DE genes set to the proportion between these two cell types. |
| ZINB-WaVE | This method takes cell types label into consideration in the parameter estimation step, thus estimation and simulation were performed directly on the entire dataset with cell type labels provided. | Estimation and simulation were performed directly on the entire dataset with cell type labels provided. We then evaluated the biological signals between the two largest cell types. |
| SparseDC | This method requires two conditions such as treatment and control, with multiple cell types in each condition, as an internal clustering step is performed to differentiate the cell types. We followed the procedure in the SparseDC documentation and split half of the cell types into condition 1 and half of the cell types into condition 2, and specified the number of clusters to be the number of cell types in condition 1 and 2. | Due to the unique setting, we did not evaluate this method for biological signals. |
| cscGAN | This method takes cell types label into consideration in the parameter estimation step, thus estimation and simulation were performed directly on the entire dataset with cell type labels provided. | Estimation and simulation were performed directly on the entire dataset with cell type labels provided. We then evaluated the biological signals between the two largest cell types. |
| * We used the procedure described in "Evaluation of biological signals" of the Methods section to calculate the proportion of differential expressed genes between the two largest cell types in the real data. This proportion was then used as the input parameter in the simulation function to control the proportion generated in the simulation data. | | |
